# Stool antigen-based enzyme immunoassays: Performance evaluations and automation

**DOI:** 10.64898/2026.01.05.697665

**Authors:** Jian R. Bao, Robert S. Jones

## Abstract

Three enzyme immunoassays (EIAs) were evaluated on two platforms each for detecting *Cryptosporidium*, *Giardia*, and *Campylobacter* in stool samples compared to gold standard methods. The *Cryptosporidium* EIAs run on stool specimens in three preservative media showed 100% agreement with direct fluorescent antibody (DFA) testing and high correlation with the microscopic method. The *Giardia* EIAs demonstrated 100% sensitivity but only moderate correlation with the microscopic method. The *Campylobacter* EIAs had 93.5% sensitivity (out of 65 positives) and 100% specificity compared to the culture method.

Automation on a DS2 system for these three EIAs, along with five others, yielded acceptable performance (92.5-100% accuracy) compared to manual methods. The automation saved labor time and improved operational efficiency but may not be cost-effective for low-volume runs due to the labor required for automation not scaling proportionally with sample numbers.

In conclusion, EIAs are preferred for detecting protozoan parasites in stool, with the *Cryptosporidium* EIA showing potential as a semi-quantitative assay and a reference method. Automation benefits high-throughput laboratories but may not be as advantageous for low-volume laboratories.

**IMPORTANCE:** Enzyme immunoassay (EIA) is a widely used method in clinical laboratories to detect pathogens in stool samples related to diarrhea diseases. This study evaluated the performance of three EIAs for detecting *Cryptosporidium*, *Giardia*, and *Campylobacter* antigens compared to their gold standard methods. The *Cryptosporidium* EIAs matched the direct fluorescent antibody (DFA) testing and had a high correlation with microscopic findings (99.7%). The *Giardia* EIAs showed 100% sensitivity but lower specificity (58%) and moderate correlation with microscopic results (87.4%). The *Campylobacter* EIA had 93.5% sensitivity (n=65 positive samples) with few discrepancies. Automating these three and five other EIAs using a DS2 system (Dynex) yielded good accuracy (92.5-100%) and 100% precision compared to manual methods. While automation saved hands-on time for high-volume assays, it may not be cost-effective for low-volume laboratories.

---

Stool antigen-based enzyme immunoassay (EIA) remains crucial in clinical microbiology for detecting diarrhea-causing pathogens. Diarrheal infections can be caused by various pathogens, such as viruses, bacteria, and parasites, leading to approximately 179 million cases of acute diarrhea in the US alone each year (1). Traditional laboratory diagnostic methods for these diseases involve manual technologies to detect causal agents in stool specimens, including microscopic observation for parasites or culture for bacteria. These techniques, which have been successful for decades, are still widely used and considered the gold standard methods (2). However, they are labor intensive, have slow turnaround times (TAT), require expertise, and may lack sensitivity and specificity. EIA-based methods offer a quicker, simpler, and more accurate alternative, making them essential in modern laboratory diagnosis (2,3,4,5).

Culture-free diagnostic methods, including EIAs, molecular assays, single-cell analysis, and artificial intelligence (AI) imaging models, are gaining popularity due to their efficiency, quick TAT, and ability to produce quality results (2,3,6). EIAs can be more sensitive and specific than microscopic methods in diagnosing intestinal protozoa (3,7) and can detect more species causing campylobacteriosis (8). Some EIA or molecular methods, such as direct fluorescent antibody (DFA) for *Cryptosporidium* (9) and nucleic acid amplification tests for viral or bacterial pathogens are considered gold standards in laboratory diagnostics (10).

EIAs use enzymes bound to antigens or antibodies to detect microbial antigens, evolving from radioimmunoassay (11). They are versatile and widely available for detecting a range of pathogens directly from stool specimens, making them ideal for rapidly identifying diverse causal agents of diarrhea. Stool in preservative media is commonly used in laboratory testing, especially for *Cryptosporidium* detection due to intermittent oocyst shedding. Preserved stool specimens, good for EIAs, may limit bacterial culture methods or cause PCR inhibitions (9). Studies of EIA performance for laboratory diagnosis with preserved stools in different preserved media and comparison to their gold standards are still scattered. In this study, we evaluated the performance of three EIAs on two platforms each for detecting enteric-pathogen-related antigens (*Cryptosporidium*, *Giardia*, and *Campylobacter*) in preserved stool compared to their respective gold standard methods. The EIAs, along with five others, were automated on a DS2 system and assessed for their performance and operational characteristics in a high-throughput laboratory setting.

## MATERIALS AND METHODS

### EIA evaluation scopes

Since 2022, three specific EIAs, along with five others, have been evaluated using two different platforms to detect some major diarrhea-causing enteric pathogens: *Cryptosporidium* spp., *Giardia* spp., and *Campylobacter* spp. from stool specimens (Table 1). *Cryptosporidium* EIAs were compared to DFA and microscopic methods, *Giardia* EIAs were compared to a microscopic method, and *Campylobacter* EIAs were compared to a culture method. Two EIA platforms were used for each of the three EIAs, including EIA kits from TechLab (Blacksburg, VA), Premier^®^ kits from Meridian Bioscience (Cincinnati, OH), or Remel ProSpecT for *Giardia* (ThermoFisher Scientific). The performance of some EIAs was compared with other EIAs when the gold standard method was unattainable.

**TABLE 1.** Enzyme immunoassay evaluation scope*.

| EIA target<br>/Platform Manufacturer** | Specimen Type | EIA Accuracy Comparison (%) (n=sample number) |  |  |
| --- | --- | --- | --- | --- |
|  |  | EIA accuracy /Gold Standard | EIA to EIA ** | Automation (DS2) to Manual |
| *Cryptosporidium*<br>/TechLab&Meridian | Preserved stools (Total Fix, Cary-Blair, formalin) | 100 (n=140) /DFA | 100<br>(n=140, p=75) | 100 (n=60, p=31) |
| *Giardia* /TechLab&Remel<br>ProspecT | Preserved stools (Total Fix, formalin, Cary-Blair or PVA) | 75 /Microscopic<br>(n=84, p=55, Sensitivity=100) | 100<br>(n=84, p=55) | 93% (n=49, p=24) |
| *Campylobacter*<br>/TechLab&Meridian | Stools in Cary-Blair/C&S | 95.7 (n=94) /Culture | 96.9<br>(n=65, p=49) | 98.1 (n=52, p=23) |
| Shiga Toxin<br>/TechLab&Meridian | Unpreserved (Cary-Blair) and other preserved Stools | NA | 97.7<br>(n=45, p=24) | 100 (n=81, p=14) |
| *H. pylori*<br>/TechLab&Meridian | Unpreserved (Cary-Blair/C&S) and preserved Stools | NA | 100<br>(n=22, p=12) | 98.0 (n=150, p=75) |
| Rotavirus<br>/Meridian | Unpreserved stool or rectal swab | NA | NA | 100 (n=45, p=19) |
| *Legionella*<br>/Binax | Urine | NA | NA | 94 (n=50, p=21) |
| Lactoferrin (Qualitative)<br>/TechLab | Unpreserved Stool | NA | NA | 96.6 (n=147, p=41) |

\*P: Positive; NA: Not applicable. \*\*The two EIA platforms as listed in the Table.

The EIAs were gradually transitioned from manual to automated methods using a DS2 system (Dynex, Chantilly, VA) as described in the EIA automation section. The performance of the EIAs under automation was evaluated, and their operational characteristics were assessed for efficiency and labor time reduction compared to manual methods.

### Specimen types and their preparations

The clinical samples used in this study were remnants of various patient specimens received in the laboratory for testing within stability-time windows under their storage conditions. Stool samples were either fresh or preserved in one of three transport media: Total Fix^®^ (Medical Chemical Corp), 10% buffered formalin (prepared in the local reagent facility), or Cary-Blair C&S transport media (Para-Pak^TM^, Meridian Diagnostics, Cincinnati, OH). Positive *Campylobacter* stools were additionally collected from other Quest Diagnostics laboratories in Texas (n=28), Florida (n=7), Missouri (n=6), and Illinois (n=2). A *Cryptosporidium* validation panel (n=20) provided by the EIA manufacturer (TechLab) was included in the study. Spiked samples were used to cover Shiga toxin-producing *E. coli*, as clinical positive specimens were rare. They were produced from negative patient samples seeded with either positive patient samples with high EIA optical density (OD) readings or liquid cultures of well-characterized organisms from CDC (AR strains, https://wwwn.cdc.gov/ARIsolateBank).

### Microscopic and culture methods

The O&P microscopic procedures followed standard laboratory protocols that were based on CDC guidelines (9). Wet mount slides were prepared from unstained or Trichrome-stained concentrated stool sediments. *Cryptosporidium* oocysts were microscopically examined using a modified acid-fast staining procedure. Prepared slides were screened at 10 x objective magnification and then at 40 x objective magnification for parasitic organism identification. Stool samples were blindly examined alongside EIAs or post EIA testing.

For *Campylobacter* culture, stool specimens were inoculated onto CAMP CVA plates (Remel, Lenexa, Kansas) and incubated at 41-43°C in a GasPak^TM^ EZ incubation container (BD, Franklin Lakes, NJ) for 24-72 hours. Isolated organisms were identified based on morphology, biochemical characteristics, and mass-spectrum method (MALDI-TOF, Bruker).

### EIA automation

The automation was implemented to enhance operational efficiency of assays while maintaining their performance. A compact DS2 fully automated ELISA system (DS2, Dynex) was utilized for EIA automation. The assay programs on the DS2 system were either provided by the EIA manufacturer (LabTech, VA) or created by the laboratory for non TechLab EIAs. Each program included all necessary steps and parameters identical to manual methods, such as liquid handling, washing, incubation, OD measurements, and data interpretation for final results. The information technology (IT) interface integrated into the DS2 system facilitated the approving process and the release of result batches from the laboratory to clinicians. EIA performance under automation was assessed by comparing to manual methods.

To adapt EIA automation, two main modifications were made for sample preparation and machine setup. Stool samples collected in testing tubes were increased in volume or mass by 2-3 times compared to the manual method. The samples were diluted with a proportionally increased diluent volume to maintain the required dilution ratios for each EIA (Table 4). The diluted stool samples were mixed thoroughly and centrifuged at 5000 x *g* for 10 minutes to precipitate solid particles. This sample preparation ensured that the sample volume was adequate for the DS2 to handle liquids smoothly and prevented potential pipette tip clogs during the assay. Barcode-labeled sample-containing testing tubes (0.5 x 5.0 mL) were scanned into the DS2 system for assay automation. The machine setup and post-run cleaning procedures are detailed in the results section.

## RESULTS

### *Cryptosporidium* EIAs compared to DFA and microscopic method

In the study involving 140 samples, 116 were clinical stool samples with 65 testing positive for *Cryptosporidium*. The remaining 24 samples were from a validation panel or survey samples (Table 2). The clinical specimens were preserved in Total-Fix (n=80, with 50 positives), Cary-Blair (n=20), or 10% Formalin (n=16).

**TABLE 2.** Performances of *Cryptosporidium* EIA for stool samples in different preservative media*.

| Stool in preservative media or origin | Total samples | *Cryptosporidium* (+) or (-) | No. of (+) or (-) sample under testing method |  |  | DFA vs EIA agreement (%) |
| --- | --- | --- | --- | --- | --- | --- |
|  |  |  | Microscopic (n=116) | DFA (n=140) | EIA ** (n=140) |  |
| Total Fix | 80 | + | 44 | 50 | 50 | 100 |
|  |  | - | 36 | 30 | 30 | 100 |
| 10% Buffered formalin | 16 | + | 5 | 5 | 5 | 100 |
|  |  | - | 11 | 11 | 11 | 100 |
| Cary-Blair/C&S | 20 | + | 10 | 10 | 10 | 100 |
|  |  | - | 10 | 10 | 10 | 100 |
| Subtotal (clinical) | 116 |  | 59+/57- | 65+/51- | 65+/51- | 100 |
| Graded survey (CAP) | 4 | + | NA | 2 | 2 | 100 |
|  |  | - | NA | 2 | 2 | 100 |
| Validation Panel (TechLab) | 20 | + | NA | 8 | 8 | 100 |
|  |  | - | NA | 12 | 12 | 100 |
| Total | 140 |  | 59+/57- | 75+/65- | 75+/65- | 100 |

\*"+", positive; "- ", negative; DFA, direct fluorescent antibody; EIA, enzyme immunoassay; NA, not applicable. \*\* EIA platforms: Meridian and TechLab have the same results.

Both EIA platforms used to detect *Cryptosporidium* spp. from the 140 samples produced identical results, showing 100% sensitivity and specificity compared to the DFA method, which is considered the gold standard (9). Stool preserved in Total-Fix^®^ was not an FDA-cleared specimen type for the TechLab EIA platform, but this study found that all three preservative stool specimen types yielded the same results as the reference DFA method for *Cryptosporidium* detection.

Among the 65 clinical *Cryptosporidium* positives identified by both DFA and EIAs, the microscopic method detected 59 (90.8%) as positives and 6 as negatives. Among the 75 negatives identified by both DFA and EIAs, the microscopic method detected 75 (100%) as negatives. The receiver operating characteristic (ROC) curve analysis showed a high correlation between the performance of the two methods (Fig. 1), with the EIA scoring 99.7% of the area under the curve (AUC) compared to the microscopic method. According to the ROC analysis, the OD cutoff value for detecting *Cryptosporidium* using the microscopic method was 0.259 (450/620 nm) from the EIA (TechLab) (Supp Table S1). The OD threshold for reliable microscopic detection of *Cryptosporidium* positive stools was 0.695. The differences in cutoff OD values between the EIA (>=0.09 for positive) and microscopic methods explain the variations in detection sensitivity.

**FIG 1.**
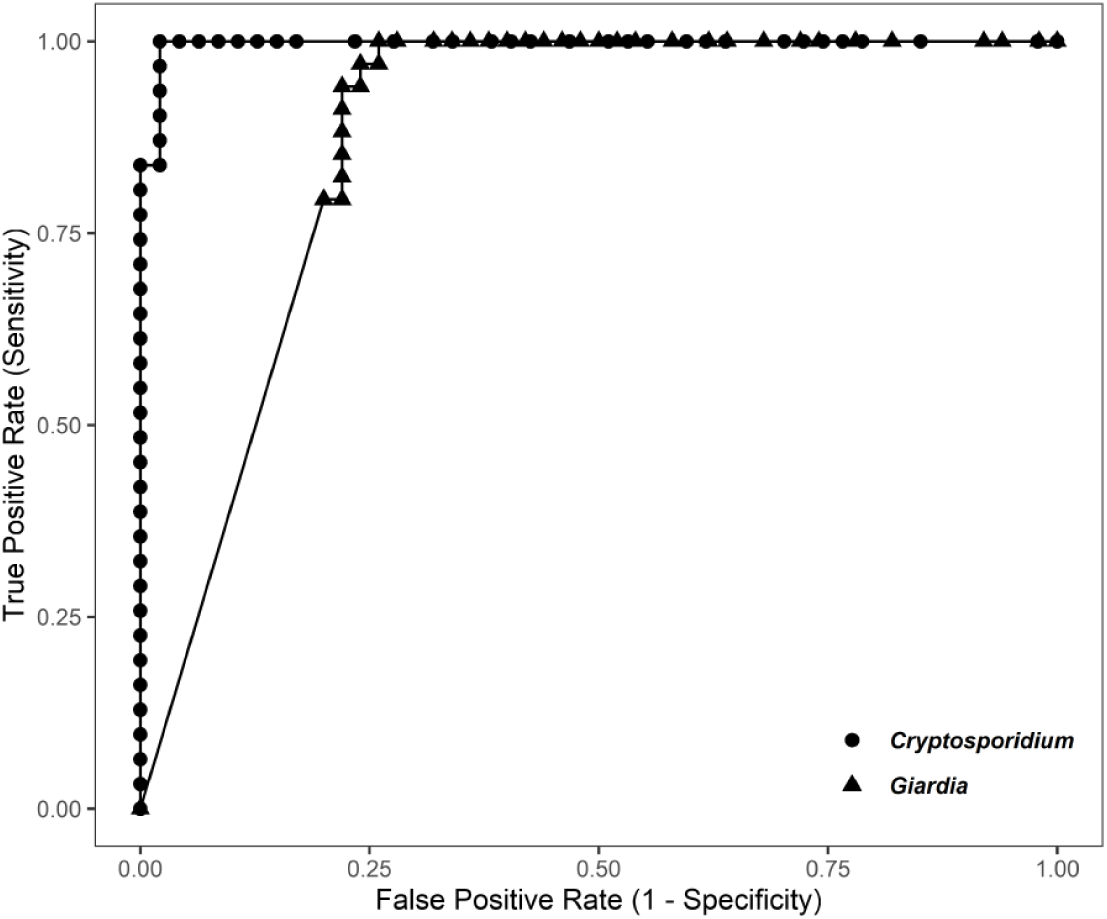
Receiver operating characteristic (ROC) curve of microscopic method compared to EIA method for detection of *Cryptosporidium* (⚫Area under curve (AUC) =0.997, n=78) or *Giardia* (▲AUC=0.874, n=50) from stool specimens.

### *Giardia* EIA and wet-mounted microscopic method

The two EIA platforms produced consistent results for the 84 stool samples, with 55 testing positive and 29 testing negative for *Giardia* antigen (Supplemental Table S2). The microscopic method identified 34 samples as *Giardia* positive and 50 as negative. All 34 samples identified as positive by the microscopic method were also positive in the EIA tests, resulting in a sensitivity of 100% for EIAs compared to 61.8% for the microscopic method. Additionally, all samples that tested negative in the EIA tests were also negative in the microscopic method. However, the microscopic method classified 21 EIA-*Giardia* positive samples as negative, resulting in a likely 58% error rate. Overall, the agreement rate between the two methods was 75%.

The ROC curve analysis showed a performance difference between EIA and the microscopic method for *Giardia* detection (Figure 1). The moderate correlation (AUC=0.874) made it difficult to establish thresholds for microscopic *Giardia* detectability based on EIA OD values. Stool samples classified as *Giardia* negative by the microscopic method exhibited a wide range of OD readings, with four samples even reaching maximum EIA OD readings (>3). Although the clinical outcomes of individuals with discrepancies in test results were not tracked in this study, the ROC curve analysis indicated that microscopic results were less consistent in detecting *Giardia* compared to EIAs.

### *Campylobacter* EIA compared to culture method

Ninety-four (94) clinical stool specimens were tested for the detection of *Campylobacter* spp. (Table 3). The culture method identified 65 *Campylobacter* positives and 29 negatives. The majority of the positive samples were identified as *C. jejuni* (n=58, 89.2%), with few as *C. coli* (n=5, 7.7%), *C. upsaliensis* (n=1, 1.5%) and one intractable isolate (1.5%).

**TABLE 3.**
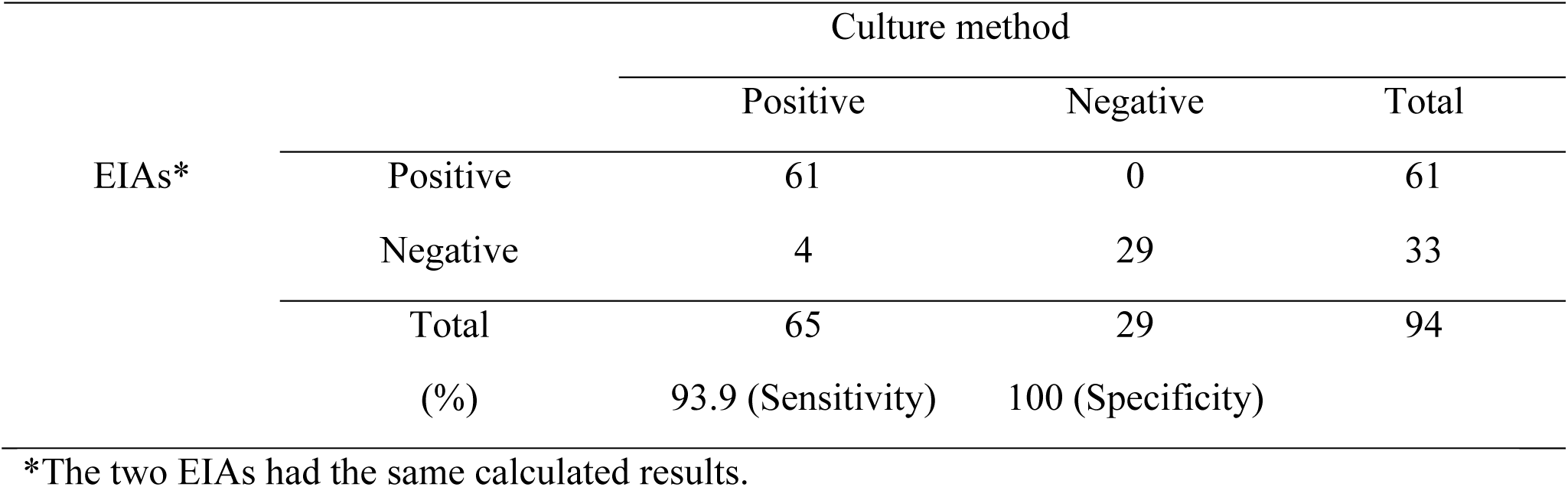
Two *Campylobacter* EIAs compared to culture method.

Both EIA platforms (Premier, Meridian and Campy CHEK, TechLab) detected 61 positives, all from culture positives (93.9% sensitivity), and 33 EIA negatives. All the culture negatives were also EIA negatives for both platforms (100% specificity). The EIAs had 100% positive predictive value (95% CI=94.48-100.00%), but their negative predictive value was undeterminable due to the selective sample collection. The culture method was more sensitive than the EIAs in detecting *Campylobacter* spp. from stools.

Each EIA platform missed four *Campylobacter* culture-positive samples as EIA negatives, with discrepancies involving five stool samples for both the EIAs (Supplement Table S3). Three culture-positive stools were identified by both EIAs as negatives and two showed conflicting results between the two platforms, being called positive by one but negative by the other EIA. PCR testing (Verigene Enteric Pathogens Nucleic Acid Test Panel, DiaSorin, Stillwater, MN) for the three culture-positive but EIA-negative samples were also *Campylobacter* negative. *C. jejuni* was not the dominant species (25%) among the culture-EIA discrepancy stools.

### EIA automation and their operational characteristics

The three EIAs mentioned earlier, along with five others, were gradually transitioned from manual to automated procedures on a DS2 system over two years (Tables 1, 4, Supplemental Table S4). The transition involved fine-tuning each assay’s performance, operational adjustments like personnel training, workflow changes, and adapting IT for data reporting.

**TABLE 4.** EIA sample processing for automation on DS2*.

| EIA | Stool specimen type | Specimen volume (µL) | Diluent volume (µL) | Centrifugation (5000 x g / 10 min) |
| --- | --- | --- | --- | --- |
| *Cryptosporidium* | preserved | 750 | NA | Yes |
| *Giardia* | preserved | 500 | NA | Yes |
|  | unpreserved | 200 or 0.2 g | 400 | Yes |
| *Campylobacter* | preserved (CB) | 200 | 200 | Yes |
|  | unpreserved | 100 or 0.1 g | 400 | Yes |
| Shiga Toxin | preserved (CB) | 200 | 200 | Yes |
|  | unpreserved | 100 or 0.1 g | 400 | Yes |
| *H. pylori* | preserved | 200 | 200 | Yes |
|  | unpreserved | 150 or 0.2 g | 600 | Yes |
| Rotavirus | stool or swab | 1st pipette mark or whole swab | 1000 | Yes |
| LUA | urine | 500, or use specimen tube directly | NA | No |
| Lactoferrin (qualitative) | stool | 50 or 0.05 g | 950 | Yes |

\*LUA: Legionella urinary antigen. The preserved specimens included Cary-Blair (CB), Total-Fix, or formalin (10%) depending on each assay's requirements. NA: not applicable.

The automated EIAs demonstrated high levels of accuracy (92.5-100%) and precision (100%) compared to their manual methods (Table 4). These results indicated that automation did not compromise EIA performance. The modifications made for automation, particularly in sample preparations, were satisfactory for stools performed on the DS2 system to produce acceptable diagnostic results. The automation in this study was run on different types of samples, from urine to stool, fresh or preserved, and produced acceptable results, indicating that the automation system and its method were suitable for various EIAs with different specimen types. Our experience with the sample preparation for automation indicated that a minimum sample volume of 400 µL per machine-fit tube (0.5 x 5.0 mL) was necessary for smooth EIA operation on the system without causing major pipetting clogs.

Using EIA kits from the same manufacturer allowed sharing common reagents (such as diluent and washing buffer, per TechLab), enabling the operation of two EIAs in a single run on the same machine. EIAs of *Cryptosporidium* and *Giardia* mostly shared the same stool samples, and their running in a single run improved operational efficiency and saved reagents, especially when sample numbers were low for both assays. While automation consumed more reagent volumes, reagent shortages were not an issue with regular EIA kits designed for manual methods. Utilizing screw-capped reagent tubes of varied sizes on the DS2 machine helped save and refrigerate left over reagents for future runs.

The automation with DS2 reduced labor time by approximately 30 minutes per 96-well EIA plate assay (Fig. 2). Efficient IT networking also decreased data processing time and result release by 10 minutes for a full run, improving workflow efficiency. While automation saved time in assay procedures, additional time was required for sample preparation and machine setup (Fig. 2). The labor times for sample centrifugation and machine setup remained consistent with minor variation based on the number of samples. This made it more labor intensive per sample for assays with fewer samples and less cost-effective for low sample volume assays or non-high-throughput laboratories.

**FIG 2.**
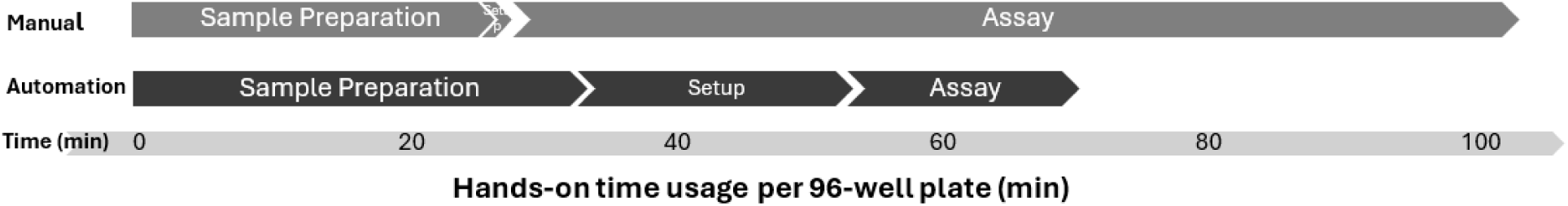
Hands-on time (min) for EIA automation on a DS2 system compared to manual EIA method.

## DISCUSSION

Rapid, sensitive, and specific diagnostic methods are crucial for managing diarrhea-related infections caused by various agents. While different laboratory diagnostic methods are available, immunoassay-based methods, particularly antigen-based EIAs, are commonly used to detect or screen the etiologies in stool samples (12). Despite concerns about specificity due to potential cross-reactions with closely related parasitic antigens and sensitivity compared to genotypic assays, EIAs have shown satisfactory performance in laboratory diagnostic practices and outweigh the potential weaknesses (13,14). In our study, both EIA platforms used to detect *Cryptosporidium* in stools showed identical results (100% sensitivity and specificity) compared to the DFA method, which is considered the gold standard (9). No cross reactions were observed between *Cryptosporidium* and *Giardia*. However, the two EIAs for *Campylobacter* had a sensitivity of 93.9% compared to the culture method. Similarly, two *Giardia* EIAs had consistent results, while the microscopic method had 70% accuracy compared to EIAs. Overall, EIAs demonstrated better performance than the microscopic method (3,7). Our results indicated that the strength of EIAs in performance depended on the targeted organisms and reference methods.

The strong correlation between *Cryptosporidium* EIA OD readings and the microscopic method (AUC=0.997, Fig. 1) indicates that EIA could be used as a semi-quantitative assay. This correlation opens the possibility of using EIA to aid in pathogen load-based diagnosis as the pathogen load is related to disease severity (15,16). The reliability of EIAs in detecting *Cryptosporidium* and *Giardia* indicates their potential as standard reference methods for identifying these parasites in stool samples (17). Various stool sample preservation methods yielded results for *Cryptosporidium* EIAs that were consistent with those of DFA (Table 2), including *Cryptosporidium* positive stool samples preserved in polyvinyl alcohol (PVA, data not shown). This suggests that different preservatives are suitable for EIAs (Table 4). The suboptimal correlation between *Giardia* EIA and the microscopic method based on ROC curve analysis (Fig. 1) may be attributed to factors such as trophozoite degradation and differences in cyst size (10-15 µm) compared to oocyst size (4-6 µm). The size differences may favor the filtration efficiency for oocysts (18), resulting in the differences of size-density ratios in prepared stool sample sediments, which made more consistent results for detecting oocysts in stool samples.

While EIA methods for detecting bacterial pathogens have received mixed reviews in terms of sensitivity and specificity (19,20,21), our study found that EIAs did not surpass the sensitivity of the culture method for *Campylobacter* detection. The PCR panel used to resolve the discrepant samples sided with EIA method results as negative for *Campylobacter* for the three culture-positive stools (Supplemental Table S3). Our results showed that C. *jejuni* was the predominant species in the culture-positive stools, accounting for 89.2% of the cases. Interestingly, *C. jejuni* was less commonly found in discrepant samples, with the presence of 25% or less (Supplemental Table S3), compared to the overall 89.2% presence. This suggests that different species may have an impact on EIA performance (22). While we did not intend to definitively claim the disproportionate presence of *C. jejuni* based on the limited number of samples in this study, it may be worth further investigation in future studies.

Automation of EIAs using the DS2 system showed promise in saving labor time, improving efficiency, and ultimately enhancing patient care. The EIA performances under automation had no significant discrepancies observed for the qualitative assays. However, the transition to automation required procedural modifications and monitoring to ensure optimal performance. Different EIA kits from the same manufacturer allowed us to share the common reagents, such as diluent and washing buffer, and thus to conduct two EIAs in a single run. An example was to run both *Cryptosporidium* EIA and *Giardia* EIA in a single run, as the two assays usually share stool samples. This approach became helpful when each assay did not have full-capacity sample numbers.

This study effectively demonstrated the competent capabilities of EIAs in laboratory diagnosis, but it could not retrieve the clinical information of the specimens examined to reveal their relationship with laboratory results. This limitation was viewed as the main drawback of the study. Although the specimens were collected from a wide range of geographic origins, the testing was conducted in a single laboratory, which could be another weakness of the study. The automation for EIAs served a general evaluation, and the performance and characteristics may vary depending on different automation systems and laboratory operation conditions. Overall, the study provided clear and useful information by evaluating different EIAs for their distinct performances under diverse conditions with sample types, platforms and procedural changes.

In conclusion, our study highlighted the performance of EIAs for detecting diarrhea-causing pathogens and their transition to automation. EIA performance varied based on the target organisms and reference methods, with *Cryptosporidium* and *Giardia* EIAs showing promising results for reference methods. *Campylobacter* EIAs had lower sensitivity than the culture method. The successful transition of EIAs to automation demonstrated potential benefits for high-throughput laboratories, with considerations for sample volume and assay type.

## ACKNOWLEDGMENTS

We thank Genaro Rubio, Giovanna Gonzales, Shahnaz Labbafi, Nam Anh Nguyen, George Deegan, Chrisopher Rayala, and Michael Gasalao for their technical assistance. We also thank other Quest Diagnostic laboratories mentioned in the paper that helped with sample collection. Special thanks to those who worked with us and contributed to achieving our goals, including Drs Travis Price, Kileen Shier, Salah Jung, Caixia Bi, and Drs Andrew Hellman and Ann Salm for their critical comments on the manuscript.

## REFERENCES

1. DuPont HL. 2014. Acute infectious diarrhea in immunocompetent adults. The New England Journal of Medicine.370:1532–1540. 10.1056/NEJMra1301069.

2. Sood N, Carbell G, Greenwald HS, Friedenberg FK. 2022. Is the medium still the message? Culture-independent diagnosis of gastrointestinal infections. Dig Dis Sci. 67:16–25. 10.1007/s10620-021-07330-6.

3. Fitri LE, Candradikusuma D, Setia YD, Wibawa PA, Iskandar A, Winaris N, Pawestri AR. 2022. Diagnostic methods of common intestinal protozoa: Current and future immunological and molecular methods. Trop Med Infect Dis. 7:253. 10.3390/tropicalmed7100253.

4. McHardy IH, Wu M, Shimizu-Cohen R, Couturier MR, Humphries RM. 2014. Detection of intestinal protozoa in the clinical laboratory. J Clin Microbiol. 52:712–20. 10.1128/JCM.02877-13.

5. Johnston SP, Ballard MM, Beach MJ, Causer L, Wilkins PP. 2003. Evaluation of three commercial assays for detection of *Giardia* and *Cryptosporidium* organisms in fecal specimens. J Clin Microbiol 41:623– 626. 10.1128/JCM.41.2.623-626.2003.

6. Zangiabadian M, Ghorbani A, Nojookambari NY, Ahmadbeigi Y, Hosseini SS, Karimi-Yazdi M, Goudarzi M, Chirani AS, Nasiri MJ. 2022. Accuracy of diagnostic assays for the detection of *Clostridioides difficile*: A systematic review and meta-analysis. J Microbiol Methods. 204:106657. 10.1016/j.mimet.2022.106657.

7. Singh A, Houpt E, Petri WA. 2009. Rapid diagnosis of intestinal parasitic protozoa, with a focus on entamoeba histolytica. Interdiscip Perspect Infect Dis. 2009:547090. 10.1155/2009/547090.

8. Buss JE, Cresse M, Doyle S. et al. 2019. *Campylobacter* culture fails to correctly detect *Campylobacter* in 30% of positive patient stool specimens compared to non-cultural methods. Eur J Clin Microbiol Infect Dis. 38:1087–1093. 10.1007/s10096-019-03499-x.

9. CDC, 2024. Laboratory diagnosis of cryptosporidiosis, Cryptosporidium spp. crypto_benchaid.pub (cdc.gov).

10. Bloomfield MG, Balm MND, Blackmore TK. 2015. Molecular testing for viral and bacterial enteric pathogens: gold standard for viruses, but don’t let culture go just yet? Pathology. 47:227–233. 10.1097/PAT.0000000000000233.

11. Engvall E, Perlmann P. 1972. Enzyme-linked immunosorbent assay, Elisa. 3. Quantitation of specific antibodies by enzyme-labeled anti-immunoglobulin in antigen-coated tubes. J Immunol. 109:129–35. 10.4049/jimmunol.109.1.129.

12. O’Leary JK, Sleator RD, Lucey B. 2021. Cryptosporidium spp. diagnosis and research in the 21st century. Food and Waterborne Parasitology. e00131. 10.1016/j.fawpar.2021.e00131.

13. Cardos AI, Maghiar A, Zaha DC, Pop O, Fritea L, Miere Groza F, Cavalu S. 2022. Evolution of diagnostic methods for *Helicobacter pylori* infections: from traditional tests to high technology, advanced sensitivity and discrimination tools. Diagnostics (Basel). 16.12:508. 10.3390/diagnostics12020508.

14. Destura R, Cena RB, Galarion MJH. et al. 2015. Advancing *Cryptosporidium* diagnostics from bench to bedside. Curr Trop Med Rep 2:150–160. 10.1007/s40475-015-0055-x)

15. Lequin RM. 2005. Enzyme immunoassay (EIA)/enzyme-linked immunosorbent assay (ELISA). Clin Chem. 51:2415–8. 10.1373/clinchem.2005.051532.

16. Platts-Mills JA, Liu J, Houpt ER. 2013. New concepts in diagnostics for infectious diarrhea. Mucosal Immunol. 6:876–85. 10.1038/mi.2013.50.

17. Mergen K, Espina N, Teal A, Madison-Antenucci S. 2020. Detecting *Cryptosporidium* in stool samples submitted to a reference laboratory. Am J Trop Med Hyg. 103:421–427. 10.4269/ajtmh.19-0792.

18. Ferguson C, Kaucner C, Krogh M, Deere D, Warnecke M. 2004. Comparison of methods for the concentration of *Cryptosporidium* oocysts and *Giardia* cysts from raw waters. Can J Microbiol. 50:675–82. 10.1139/w04-059. PMID: 15644920.

19. Bessède E, Delcamp A, Sifré E, Buissonnière A, Mégraud F. 2011. New Methods for detection of Campylobacters in stool samples in comparison to culture. J Clin Microbiol. 49.10.1128/jcm.01489-10.

20. Gerritzen A, Wittke JW, Wolff D. 2011. Rapid and sensitive detection of Shiga toxin-producing *Escherichia coli* directly from stool samples by real-time PCR in comparison to culture, enzyme immunoassay and Vero Cell cytotoxicity assay. Clin Lab. 57:993–8. PMID: 22239032.

21. Mank TG, Zaat JO, Deelder AM, van Eijk JT, Polderman AM. 1997. Sensitivity of microscopy versus enzyme immunoassay in the laboratory diagnosis of giardiasis. Eur J Clin Microbiol Infect Dis. 16:615–9. 10.1007/BF02447929.

22. Franco J, Bénejat L, Ducournau A, Mégraud F, Lehours P, Bessède E. 2021. Evaluation of CAMPYLOBACTER QUIK CHEK™ rapid membrane enzyme immunoassay to detect *Campylobacter* spp. antigen in stool samples. Gut Pathog. 13:4. 10.1186/s13099-021-00400-0.

